# Reduced frontal gamma power at 24 months is associated with better expressive language in toddlers at risk for autism

**DOI:** 10.1101/430421

**Authors:** Carol L. Wilkinson, April R. Levin, Laurel J. Gabard-Durnam, Helen Tager-Flusberg, Charles A. Nelson

**Affiliations:** Division of Developmental Medicine, Boston Children’s Hospital, 1 Autumn Street, 6th Floor, Boston, MA 02115; Department of Neurology, Boston Children’s Hospital, 300 Longwood Avenue, Fegan 11^th^ Floor, Boston, Massachusetts, 02115; Department of Psychological and Brain Sciences, Boston University, 64 Cummington Mall Boston, Massachusetts, 02215

## Abstract

Gamma oscillations have been associated with early language development in typically developing toddlers, and gamma band abnormalities have been observed in individuals with ASD, as well high-risk infant siblings (those having an older sibling with autism), as early as 6-months of age. The current study investigated differences in baseline frontal gamma power and its association with language development in toddlers at high versus low familial risk for autism. EEG recordings as well as cognitive and behavioral assessments were acquired at 24-months as part of prospective, longitudinal study of infant siblings of children with and without autism. Diagnosis of autism was determined at 24–36 months, and data was analyzed across three outcome groups - low risk without ASD (n=43), high-risk without ASD (n=42), and high-risk with ASD (n=16). High-risk toddlers *without* ASD had reduced baseline frontal gamma power (30–50Hz) compared to low-risk toddlers. Among high-risk toddlers increased frontal gamma was only marginally associated with ASD diagnosis (p=0.06), but significantly associated with reduced expressive language ability (p=0.007). No association between gamma power and language was present in the low-risk group. These findings suggest that differences in gamma oscillations in high-risk toddlers may represent compensatory mechanisms associated with improved developmental outcomes.

## INTRODUCTION

Autism spectrum disorder (ASD) is defined by (1) deficits in social communication or interaction, and (2) restricted or repetitive behaviors(1). However individuals with ASD are remarkably heterogeneous in their phenotype - both in the presentation of core symptoms, as well as associated key developmental milestones such as language and cognitive development. Furthermore, language development of toddlers diagnosed with ASD can be quite variable, with 30% being minimally verbal by school-age, and roughly one-quarter developing age-appropriate expressive language skills(2,3). In fact, language acquisition by the end of preschool is one of the best predictors of later achievement and functioning(4–8). As such, it is important to identify early brain factors that not only influence the development of the core symptoms in ASD, but also impact language development.

A goal in improving the functional outcomes of children with autism is to identify those at greatest risk as early in life as possible, often before the behavioral repertoire of the infant is sufficiently mature to reveal consistent signs of the disorder. In this context a great deal of recent attention has been paid to recording the brain’s electrical activity using electroencephalography (EEG) from infants at high risk for developing autism by virtue of having an older sibling with the disorder(9–13). EEG measured gamma oscillations (~30–80Hz) are of particular interest in ASD as they are associated with higher order cognitive processes including sensory integration, as well as information and language processing(14–18). In addition, gamma oscillations are modulated by GABA-ergic inhibitory interneurons, which are implicated in the pathophysiology of ASD and other neurodevelopmental disorders(19–22). Many studies have reported differences in gamma-band power in older children or adults with ASD compared to individuals without ASD, however most studies do not examine correlations with clinical symptoms(23–29), making it difficult to determine whether these differences are primary causes of impairments, or the result of ongoing compensatory mechanisms. Recent work in typically developing infants also supports a role for gamma in early language acquisition and development. For example, by 6 months of age, infants display increased gamma-band activity in response to native, but not non-native speech(16). In addition, resting frontal gamma power has been associated with both receptive and expressive language ability(17,18,30). Work by Benasich et al. has found that resting frontal gamma power is reduced in toddlers aged 24 and 36 months who have a family history of language impairment, and that gamma power is positively correlated with current language ability across a combined population of toddlers with and without family history of language impairment(17). However, in teenagers, resting gamma is negatively correlated with measures of reading ability(31), suggesting that the role of resting gamma oscillations on language processes may be dependent on the age of the individual.

Longitudinal studies following infants at increased risk of ASD provide us the opportunity to further tease apart the functional significance of these neurophysiological differences. Infant siblings of children with ASD have an increased incidence of ASD diagnosis, currently estimated to be as high as 1 in 5(32). Accumulating research suggests that there are significant neurobiological differences, including gamma oscillations, in these high-risk infant siblings (as compared to siblings of typically developing children) that are present well before symptom onset, and even among high-risk infants who do not later develop ASD^11, 13, 28–35^. For example, our group reported that at both 3 and 6 months of age, high-risk infants, regardless of their later diagnosis, show reduced frontal EEG power across many frequencies(13,41). With regards to gamma oscillations, our lab using a subset of the data presented in this paper, found differences between low and high-risk groups in the baseline frontal gamma power developmental trajectory(13) - the high-risk group had lower frontal gamma power at 6 months of age, but had similar gamma power by 24 months. This previous analysis however did not separate the high-risk group by ASD outcome, and did not correlate gamma power differences with concurrent or future language measures.

In the present study we address three aims. First, using an expanded data set, we assessed whether baseline frontal gamma power at 24 months is altered between three outcome groups - low-risk without ASD (LR), high-risk without ASD (HR-NoASD), and high-risk with ASD (HR-ASD). Second, we assessed whether frontal gamma power at 24 months was associated concurrent or future language ability, and whether these brain-behavior associations were different between outcome groups. Finally, given the mounting evidence that the pathophysiology and phenotype of ASD may be different between males and females, we investigated within-group differences between sexes and present data both combined and stratified by sex.

## MATERIALS/SUBJECTS AND METHODS

### Participants

Infants were enrolled in a comprehensive longitudinal study of early neurocognitive development of infant siblings of children with ASD, conducted at Boston Children’s Hospital/Harvard Medical School and Boston University. Institutional review board approval was obtained from Boston University and Boston Children’s Hospital (#X06–08–0374) prior to starting the study. Written, informed consent was obtained from all parents or guardians prior to their children’s participation in the study.

All infants had a minimum gestational age of 36 weeks, no history of prenatal or postnatal medical or neurological problems, and no known genetic disorders (e.g., fragile-X, tuberous sclerosis). Furthermore, all infants were from primarily English-speaking households (English spoken more than 75% of the time). Infants designated as high-risk for ASD (HR) were defined by having at least one full sibling with a DSM-IV ASD diagnosis that could not be attributed to a known genetic disorder. All older siblings had a community diagnosis of ASD, and in the majority of cases this was confirmed using the Social Communication Questionnaire (SCQ)(42) and/or the Autism Diagnostic Observation Schedule (ADOS)(43). Three older siblings were under the age of 4, and the Pervasive Developmental Disorders Screening Test-II (PDDST-II)(44) was used instead of the SCQ. Five older siblings did not have an SCQ or ADOS completed, however they were all diagnosed in specialty clinics with expertise in ASD evaluation.

Low-risk infants were defined by having a typically developing older sibling and no first- or second-degree family members with ASD. In the majority of cases the siblings of LR infants were screened for ASD (67/72) using the SCQ or PDD-ST-II, followed by the ADOS if concerns of ASD were raised.

A total of 255 participants were enrolled in the study. Given the longitudinal nature of the study and enrollment at an early age, 16 participants were excluded after enrollment as additional information was gathered and children no longer met our inclusion or exclusion criteria. In addition, 3 participants were excluded due to medical reasons that occurred during the study (diagnosis of hearing impairment, seizures, and genetic finding associated with developmental delays).

Only a portion of the enrolled participants had high quality EEG recorded at the 24- month time point, and were therefore included in the analysis of this study. In addition, three low-risk males went on to meet criteria for ASD and were not included in further analysis. Ultimately 43 LR and 58 HR toddlers were included in the analysis. Of the 58 HR toddlers, 16 (27.6%) met criteria for ASD (Table 1).

**Table 1.** Demographic Information of Sample Participants

| Characteristic | LR<br>n = 43 | HR-NoASD<br>n = 42 | HR - ASD<br>n = 16 | Fisher's Exact Test<br>*p* value |
| --- | --- | --- | --- | --- |
| **Sex** | 24M, 19F | 20M, 22F | 11M, 5F | *p* = 0.36 |
| **Maternal Education, *n* (%)** |  |  |  | *p* = 0.01 |
| Not answered | 5 (11.6) | 1 (2.4) | 4 (25.0) |  |
| < 4-year college degree | 2 (4.7) | 9 (21.4) | 3 (18.8) |  |
| 4-year college degree | 8 (18.6) | 7 (16.7) | 6 (37.5) |  |
| Graduate degree | 28 (65.1) | 25 (59.5) | 3 (18.8) |  |
| **Paternal Education, *n* (%)** |  |  |  | *p* = 0.17 |
| Not answered | 5 (11.6) | 1 (2.4) | 5 (16.6) |  |
| < 4-year college degree | 3 (7.0) | 3 (7.1) | 7 (23.3) |  |
| 4-year college degree | 12 (27.9) | 15 (35.7) | 10 (33.3) |  |
| Graduate degree | 23 (53.5) | 23 (54.8) | 8 (26.7) |  |
| **Household Income** |  |  |  | 0.67 |
| Not answered | 7 (16.3) | 2 (4.8) | 5 (31.3) |  |
| <\$75,000 | 5 (11.6) | 4 (9.5) | 2 (12.5) | |
| >\$75,000 | 31 (72.1) | 36 (85.7) | 9 (56.2) | |
| **Race** |  |  |  | 0.07 |
| Non-White | 6 (14.0) | 2 (4.8) | 4 (25.0) |  |
| **Ethnicity** |  |  |  | 0.25 |
| Hispanic or Latino | 2 (4.7) | 2 (4.8) | 1 (2.3) |  |

Abbreviations: ASD, autism spectrum disorder; LR, low-risk without ASD; HR-NoASD, high-risk without ASD; HR-ASD, high-risk with ASD;

### Behavioral Assessment

The Mullen Scales of Early Learning (MSEL) were administered at 6, 12, 18, 24, and 36 months of age by trained examiners. Age-standardized T-scores in four domains (receptive language, expressive language, fine motor, and visual reception) were used to assess development. The ADOS was administered at 18, 24, and 36 months of age by research staff with extensive experience in testing children with developmental disorders, and then co-scored by an ADOS-reliable research assistant via video recording. For those children meeting criteria on the ADOS, or coming within 3 points of cutoffs, a Licensed Clinical Psychologist reviewed video recordings of behavioral assessments and scores, and provided a best estimate clinical judgment: typically developing, ASD, or non-spectrum disorder (e.g. ADHD, anxiety, language concerns). For this analysis, HR infants receiving a clinical judgment of either typically developing or non-spectrum disorders were classified as HR-NoASD, and those receiving a clinical judgment of ASD were classified as HR-ASD. All infants classified as HR-ASD were administered an ADOS at 24 and/or 36 months. Calibrated severity scores for the 24-month ADOS were determined to allow for comparison between individuals administered different ADOS modules.

### EEG Assessment

Baseline EEG data were collected at 24 months of age in a dimly lit, sound-attenuated, electrically shielded room. The infant was held by their seated caregiver during data collection while a research assistant ensured the infant remained calm and still by blowing bubbles and/or showing toys. Continuous EEG was recorded for 2–5 minutes. EEG data were collected using either a 64-channel Geodesic Sensor Net System or a 128-channel Hydrocel Geodesic Sensor Nets (Electrical Geodesics, Inc., Eugene, OR, USA) connected to a DC-coupled amplifier (Net Amps 200 or Net Amps 300, Electrical Geodesics Inc.). There was no difference in distribution of net type between outcome groups (X^2^_4_ = 1.912, *p*=0.38). Data were sampled at either 250 Hz or 500 Hz and referenced to a single vertex electrode (Cz), with impedances kept below 100kΩ. Electrooculographic electrodes were removed to improve the infant’s comfort during data collection.

### EEG pre-processing

The continuous, non-task related EEG portion of the raw NetStation (EGI, Inc, Eugene, OR) files were exported to MATLAB (versionR2017a) for pre-processing and subsequent power analysis. All files were batch processed using the Batch EEG Automated Processing Platform (BEAPP - https://github.com/lcnbeapp/beapp) to ensure uniform analysis regardless of when the EEG was acquired or which risk group they were in. A 1-Hz high-pass filter and 100Hz low-pass filter were applied. Data sampled at 500 Hz were resampled using interpolation to 250 Hz. Both experimental and participant-induced artifacts were then identified and removed using the Harvard Automated Preprocessing Pipeline for EEG (HAPPE), a MATLAB based pre-processing pipeline optimized for developmental data with short recordings and/or high levels of artifact, to automate pre-processing and artifact removal, and to evaluate data quality in the processed EEGs(45). While historically artifact removal has largely been accomplished through visual inspection, more recently the field has moved to more automated techniques that are less prone to human error and subjectivity, and allow for increased retention in data for analysis. HAPPE has been shown to both reject a greater proportion of artifact while simultaneously preserving underlying signal relative to manual editing. HAPPE also provides data output quality measures that can be used to systematically reject poor quality data unfit for further analyses. HAPPE artifact identification and removal includes removing 60Hz line noise, bad channel rejection, and participant produced artifact (eye blinks, movement, muscle activity) through wavelet-enhanced independent component analysis (ICA) and MARA (Multiple Artifact Rejection Algorithm)(46,47). MARA was, in part, chosen for its excellent detection and removal of muscle artifact components which can affect gamma signal(45,46). The following channels, in addition to the 10–20 electrodes, were used for MARA: 64-channel net - 2, 3, 8, 9, 12, 16, 21, 25, 50, 53, 57, 58; 128-channel net - 3, 4, 13, 19, 20, 23, 27, 28, 40, 41, 46, 47, 75, 98, 102, 103, 109, 112, 117, 118, 123, After artifact removal using HAPPE, data were re-referenced to an average reference. Data were then detrended using the signal mean, and then regions of high-amplitude signal (>40 uV was used to account for the reduce signal amplitude post HAPPE processing) were removed prior to segmenting the remaining data into 2-second windows to allow for power calculations using multitaper spectral analysis(48). Non-continuous data were not concatenated.

### EEG power analysis

A multitaper fast Fourier transform, using three orthogonal tapers(49) was used to calculate a power spectrum on each segment for the following frontal electrodes: 64-channel net - 2, 3, 8, 9, 12, 13, 58, 62.; 128-channel net - 3, 4, 11, 19, 20, 23, 24, 27, 118, 123, 124 (Supplemental Figure 1). For each individual EEG and each electrode, the average power across all two-second segments was then calculated for the gamma band, defined as [30–50Hz]. Gamma power was then averaged across the listed electrodes for each individual to obtain their average frontal gamma power. Here we report absolute power values, normalized by a log 10 transform.

**Figure 1.**
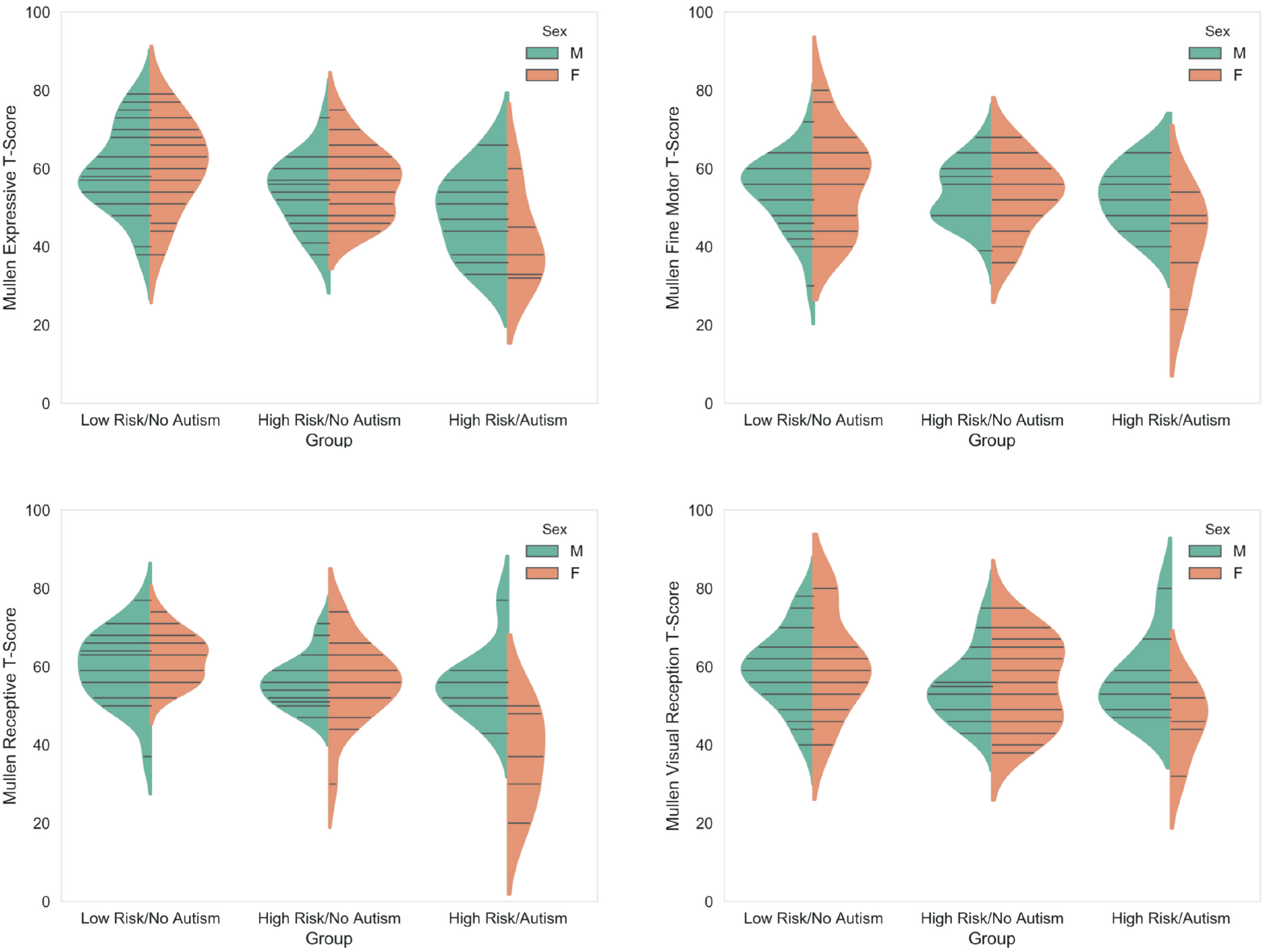
Mullen Scales of Early Learning T-Scores. Violin plots of T-scores from each of the 4 subscales (Expressive Language, Receptive Language, Fine motor, and Visual Reception) are shown for each outcome group, divided into males and females. Lines represent individual data points. LR (n: males = 24, females 19), HR-NoASD (n; males 18–19, females 20–21); HR-ASD (n: males 10, females 5).

EEG rejection criteria: EEGs were rejected if they had fewer than 20 segments (40 seconds of total EEG), or were more than 3 standard deviations from the mean on the following HAPPE data quality output parameters: percent good channels (< 82%), mean retained artifact probability (<0.3), median retained artifact probability (<0.35), percent of independent components rejected (<84%), and percent variance retained after artifact removal (>32%). Based on the above criteria, 8 of the 148 EEGs collected at 24 months were rejected. Additionally, any EEG with a mean gamma power greater or less than two SD from their outcome group mean were reviewed blind to outcome group, leading to two additional 24 month EEGs to be rejected. Furthermore, within the remaining data set, HAPPE data quality output parameters were not significantly correlated (Pearson’s r values ranged from −0.16 to 0.1) with mean frontal gamma power, supporting adequate removal of muscle artifact by HAPPE. We have also previously shown that the distribution of each of the above HAPPE data quality output parameters are similar across the three outcome groups(45).

### Statistical Analyses

In Tables 1 and 2 all categorical variables are presented as frequencies and percentages, and continuous variables are presented as means and standard deviations. A Fisher-exact test was used to characterize differences in demographic data between groups. All continuous variables within each outcome group were normally distributed using the Shapiro-Wilks test. Two-way ANOVA, followed by post-hoc Bonferroni tests for multiple comparisons, were used to determine effects of group, sex, and group x sex interactions on head circumference, MSEL scores, ADOS calibrated severity scores, and frontal gamma power.

Logistic regression was used to determine whether frontal gamma power was associated with ASD diagnosis. Multivariate linear regression was used to characterize the relationship between frontal gamma power and MSEL language scores at 24 and 36 months. Multiple comparisons within models were adjusted for using False Discovery Rate.

All reported P values are two-tailed, with a P value of 0.05 indicating statistical significance. Analyses were performed using Stata software, version 14.2 (Stata). Figures were created using Python 2.7 and python data visualization libraries (*matplotlib*(50) and *Seaborn (https://seaborn.pydata.org/index.html)*).

## RESULTS

### Sample Description

The demographic data for each outcome group (LR, HR-NoASD, and HR-ASD) are provided in Table 1. While there was no significant difference in proportion of males between groups, there were more than twice as many male HR-ASD than female HR-ASD toddlers. There was a significant group difference in maternal, but not paternal education, with a higher proportion of mothers with less than a college degree in the HR-NoASD and HR-ASD groups compared to the LR group. There were no differences in household income, race, or ethnicity. Notably, the majority of participants were white with household income above $75,000.

### Head Circumference

Given recent reports of increased head circumference and early brain overgrowth in ASD populations, we examined whether there were differences in head circumference at 24 months within our sample population. There were no differences in head circumference between groups, however there were expected differences between males and females, with females having smaller head sizes in all groups (F(1,93) = 12.68, p=.0006). There was no effect of group or group × sex interactions on head circumference (Table 2).

**Table 2.** 24-Month Sample Characteristics

| Characteristics | LR-Negative<br>mean±SD |  |  | HR-Negative |  |  | HR-ASD |  |  | p-Value |  |  |
| --- | --- | --- | --- | --- | --- | --- | --- | --- | --- | --- | --- | --- |
|  | Total | Male | Female | Total | Male | Female | Total | Male | Female | Group | Sex | GroupX<br>Sex |
| Head Circumference<sup>†</sup> | 49.3±1.5<br>(n=42) | 50.0±1.1<br>(n=24) | 48.4±1.4<br>(n=18) | 49.2±1.6<br>(n=41) | 49.8±1.8<br>(n=20) | 48.5±1.3<br>(n=21) | 49.4±1.4<br>(n=16) | 49.6±1.6<br>(n=11) | 49±0.7<br>(n=5) | df(2,93)<br>F=0.06<br>0.946 | df(1,93)<br>F=12.68<br>0.0006 | df(2,93)<br>F=0.70<br>0.499 |
| Mullen T-Scores<sup>‡</sup> | n = 43 | n = 24 | n = 19 | n = 38-39 | n = 18-19 | n = 20-21 | n = 15 | n = 10 | n = 5 | df(2,78) | df(1,78) | df(2,78) |
| Expressive Language | 60.2±10.8 | 59.5±10.8 | 61.0±11.1 | 55.2±8.8 | 54.1±8.8 | 56.2±8.9 | 46.1±11.0 | 48.3±10.6 | 41.6±11.5 | F=4.35<br>0.016 | F=0.85<br>0.36 | F=0.03<br>0.98 |
| Receptive Language | 60.8±7.8 | 59.7±9.0 | 62.2±6.0 | 56.2±8.4 | 56.1±6.4 | 56.3±10.1 | 49.5±13.4 | 55.8±8.9 | 37±12.5 | F=10.75<br>0.0001 | F=5.33<br>0.02 | F=6.26<br>0.003 |
| Fine Motor | 55.1±10.6 | 54.1±9.1 | 56.5±12.3 | 53.5±8.5 | 53.6±7.6 | 53.3±9.4 | 49.2±10.6 | 53±8.1 | 41.6±11.8 | F=1.31<br>0.28 | F=0.47<br>0.49 | F=1.06<br>0.35 |
| Visual Reception | 59.9±10.6 | 59.5±9.5 | 60.5±12.6 | 55.4±9.9 | 54.9±8.6 | 55.8±11.1 | 53.2±10.9 | 56.8±10.3 | 46±9.2 | F=1.27<br>0.29 | F=0.13<br>0.71 | F=0.79<br>0.46 |
| ADOS<sup>§</sup> | n = 36 | n = 19 | n = 14 | n = 41 | n = 20 | n = 21 | n = 16 | n = 11 | n = 5 |  |  |  |
| 24m Severity Score | 1.63±0.96 | 1.58±0.9 | 1.29±0.61 | 2.02±1.25 | 2.35±0.61 | 1.71±0.78 | 4.56±2.58 | 3.55±2.11 | 6.8±2.17 | F=30.4<br><0.0001 | F=3.35<br>0.07 | F=8.69<br>0.0004 |

Abbreviations: ASD, autism spectrum disorder; LR-Negative, low-risk without ASD; HR-Negative, high-risk without ASD; HR-ASD, high-risk with ASD; ADOS, Autism Diagnostic Observation Schedule
† Two-way ANOVA
‡ Two-way ANOVA with Maternal Education as covariate
§ ANCOVA with MSEL Expressive and Receptive Language T-scores as covariates

### Group and Sex differences in Developmental Profiles

We next examined group (LR, HR-NoASD, HR-ASD) and sex differences, as well as possible within-group sex differences on the MSEL subscales (Expressive and Receptive Language, Fine Motor, and Visual Reception). Given differences in maternal education between groups at this time point, maternal education was included in the model as a covariate. There was a significant main effect of group on Expressive Language, and a significant interaction between effects of sex and group on Receptive Language (Table 2, Figure 1). Specifically, HR-ASD toddlers had significantly lower MSEL Expressive T-scores compared to LR toddlers (p=0.02, Bonferroni). For Receptive Language, further post-hoc analyses found significant group differences for females but not males (p<0.005, Bonferroni), and that females in the HR-ASD group had lower Receptive Language T-scores compared to males (p=0.01, Bonferroni). There were no effects of group or sex, or interaction effects of group and sex, on Fine Motor or Visual Reception measures.

Next we examined sex differences in ASD symptoms at 24 months, using the ADOS severity score as the dependent variable and group, sex, and group×sex interactions as independent variables (Table 2). To control for possible confounding of language ability on ADOS severity, MSEL Expressive and Receptive Language T-Scores were included as covariates. There was a significant interaction between the effects of sex and group. Post-hoc analyses showed that both male and female HR-ASD toddlers had significantly increased severity scores compared to their respective counterparts in the LR group (p=0.006; p<0.001, Bonferroni). In addition, HR-ASD females had significantly increased severity scores compared to HR-NoASD females (p<0.001), however HR-ASD males had only marginally significant increased severity scores compared to HR-NoASDs males (p=0.06). In line with this, HR-ASD females had significantly higher ADOS severity scores compared to HR-ASD males (p=0.005).

Overall, in this study sample, high-risk females with ASD had the lowest expressive and receptive language scores, and highest ADOS severity scores. No differences between groups were observed for measures of Fine Motor and Visual Reception skills.

#### Frontal Gamma Power

Next we asked whether there were group or sex differences in baseline frontal gamma power at 24 months of age. We hypothesized that the HR-ASD group would have significantly different frontal gamma power compared to LR and HR-NoASD groups. Two-way ANOVA was used determine whether there were effects of outcome group or sex, as well as possible group×sex interactions, on mean frontal gamma power. Given differences in head circumference between sexes, and differences in maternal education between groups, both were included as covariates.

A main effect of outcome group was present (F_2,80_=4.73, p =0.01; Figure 2), however, contrary to our expectations, we found this was not due to HR-ASD differences, but rather reduced gamma power in the HR-NoASD group when compared to LR controls (p=0.013, Bonferroni). There was no difference between males and females, and no significant group×sex interactions.

**Figure 2.**
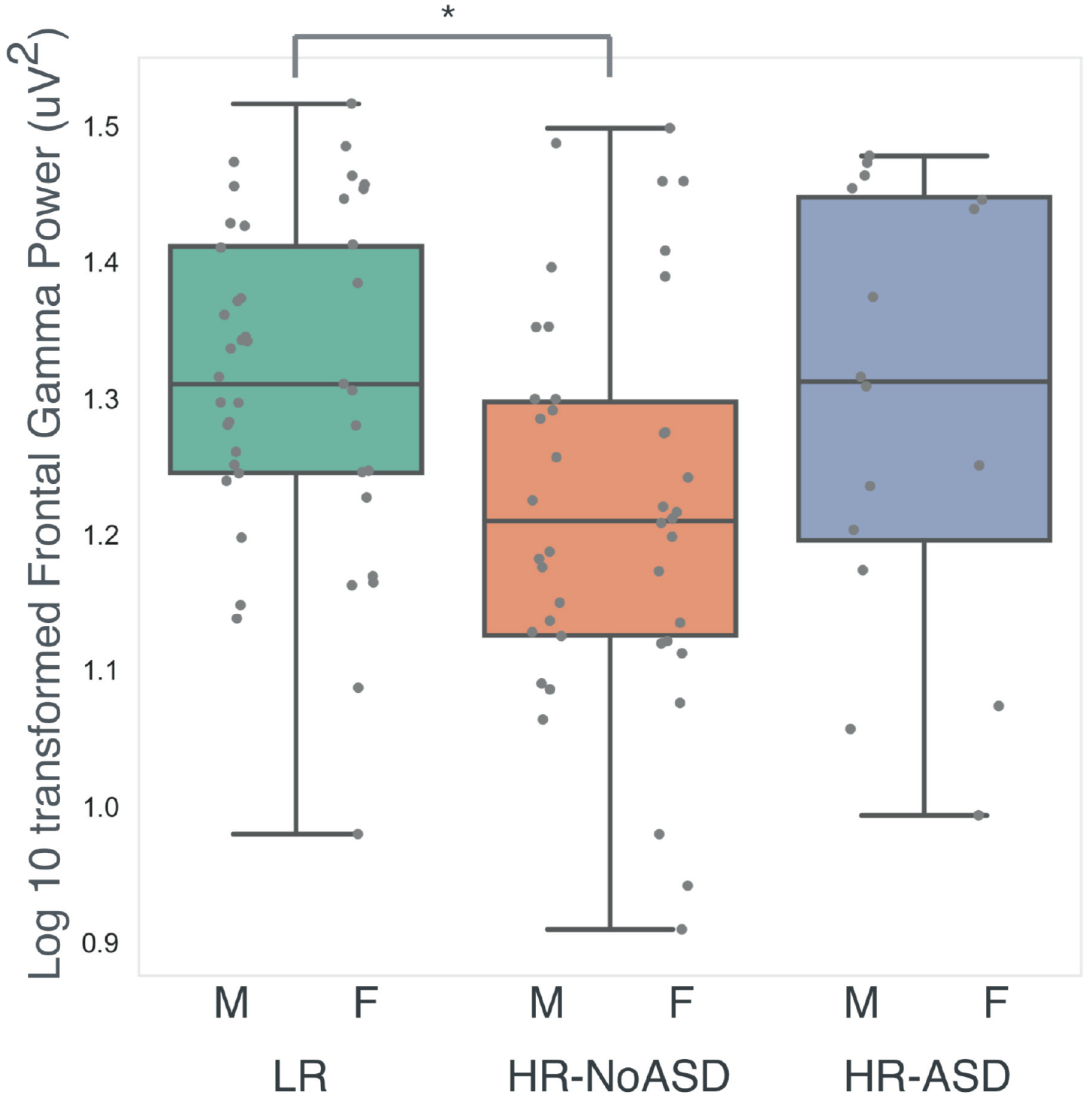
Frontal Gamma Power reduced in HR-NoASD Group. Box plots of Frontal Gamma Power are shown for each outcome group. Individual data points for males (left) and females (right) are also shown for each group. Mean values: LR (n=43, 1.31±0.12); HR-NoASD (n=42, 1.22±0.14); HR-ASD (n = 16, 1.30±0.16). Two-way ANOVA test, controlling for head circumference and maternal education, showed main effect of group (F2,80=4.73, p =0.01) with reduced frontal gamma in HR-NoASD group compared to LR group (Bonferroni, p=0.013).

This finding suggests that *within* a high-risk population, increased frontal gamma at 24 months of age may be associated with ASD diagnosis. However, within the high-risk population frontal gamma power was only marginally associated with ASD diagnosis in a logistic-regression model that adjusted for sex and maternal education (odds ratio per 1-SD increase in frontal gamma power, 2.1; 95% CI, 0.98 to 4.6, p = 0.06). In addition this association was further reduced when MSEL Verbal Quotient was added as a covariate (odds ratio, 1.5, 95% CI, 0.6 to 3.48, p = 0.4), emphasizing the strong known relationship between ASD diagnosis and language skills.

#### Frontal Gamma and Concurrent MSEL Language Scores

The close relationship between language and ASD outcome creates challenges in identifying neural correlates that are specific to ASD. Do aberrant gamma measurements in ASD populations represent brain changes that are specific to ASD, or do they represent highly associated developmental phenotypes, such as language delay or cognitive challenges, that are not core features of ASD? In the present study’s sample population, reduced frontal power across multiple frequency bands is observed at 3 months of age in the high risk group, well before ASD symptoms are present(41), and remain reduced in the HR-NoASD, but not the HR-ASD group at 24 months of age. This suggests that aberrant gamma oscillations may not be specific to ASD outcome, but a broader developmental process. To investigate this further, we next asked whether the relationships between frontal gamma power and MSEL language scores are different between risk and outcome groups.

Initially, using simple, unadjusted, Pearson correlations (Figure 3) between risk groups, we found in high-risk toddlers that frontal gamma power was negatively correlated with MSEL Expressive (r = −0.24, p=0.01, n=54), but not Receptive T-scores (r=-0.2, p=0.15, n=54). No correlation between gamma and language scores was observed in low risk toddlers (Expressive: r=0.01, p=0.94, Receptive: r=0.04, p=0.8; n=43). When the highrisk group was divided into outcome groups, this negative correlation between frontal gamma and expressive language was maintained in the HR-NoASD group (r=-0.31, p=0.05, n=39). A similar, but not significant trend was observed in the HR-ASD group.

**Figure 3.**
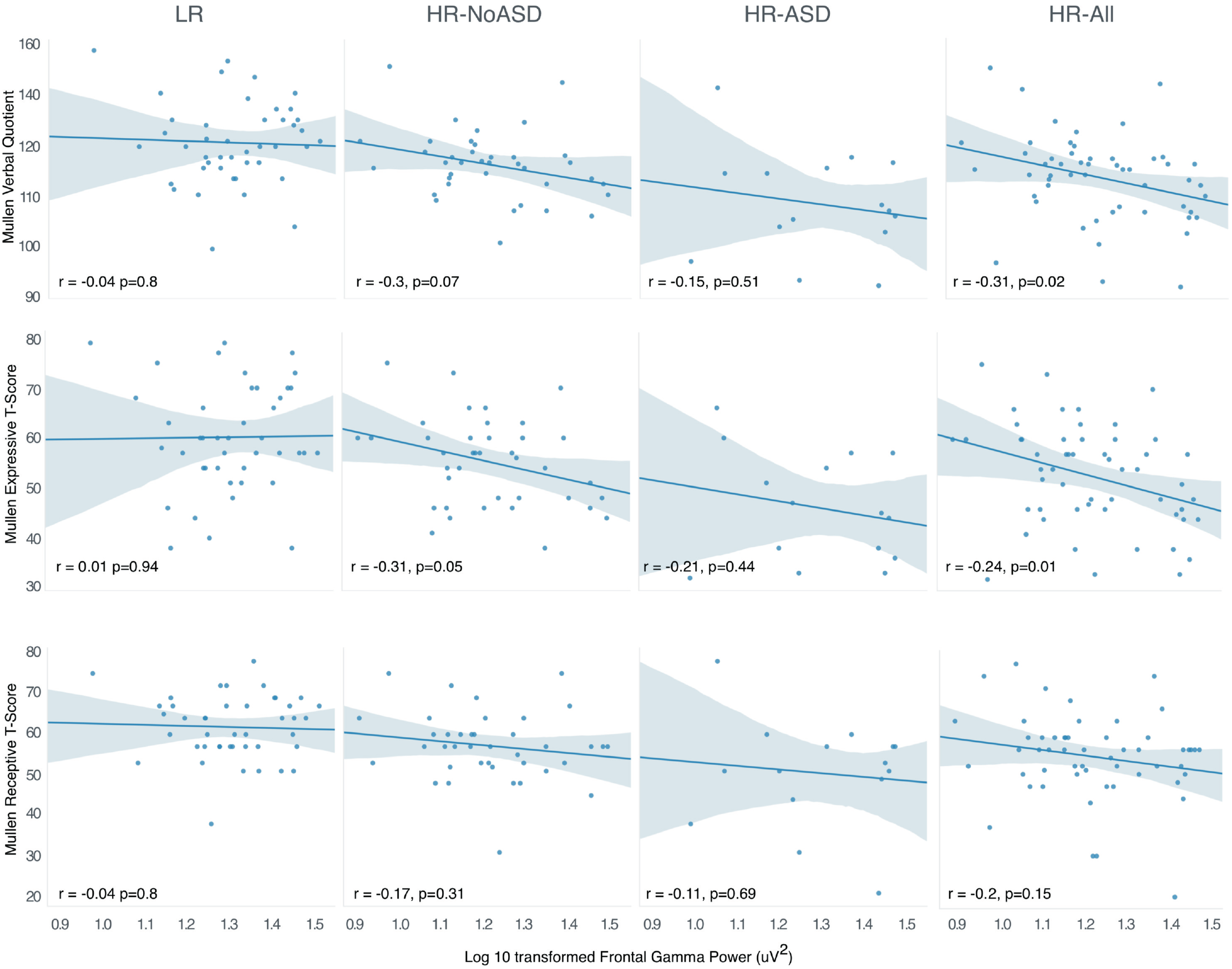
Frontal Gamma Power and Mullen Scales of Early Learning Language Scores. Frontal gamma is negatively correlated with the Mullen Verbal Quotient score for high-risk toddlers (HR-ALL), but not low-risk toddlers (LR). When divided into language subscales, this negative correlation was only significant for Expressive, but not Receptive Language T-scores. When further divided into outcome groups, only high-risk toddlers without autism (HR-NoASD) showed significant negative correlation between frontal gamma and expressive language T-scores.

In order to evaluate further the effect of risk and outcome group on the relationship between frontal power and expressive language, as well as to describe any within-group differences between males and females, two linear regression models were further examined, using MSEL Expressive T-Score as the dependent variable. Model 1 (Adjusted R^2^ = 0.16) included both two-way and three-way interactions between risk (low versus high risk), sex, and frontal gamma. Model 2 (Adjusted R^2^ = 0.17) included both two-way and three-way interactions between outcome group (LR, HR-NoASD, HR-ASD), sex, and frontal gamma. Three-way interactions for both models had p-values less than 0.25 and were therefore retained (Table 3). Both models also included head circumference and maternal education as covariates given their differences between sex and group respectively within our data set. In order to specifically evaluate the relationship between MSEL Expressive Language T-scores and frontal gamma power within risk or outcome subgroups, a marginal effects analysis was conducted and slopes are presented in Table 3.

**Table 3.** Effect of 24m Frontal Gamma Power on MSEL Expressive T-Score^†^

| Model 3-way interactions |  | P Value |
| --- | --- | --- |
| Model 1 | sex x risk x gamma | 0.137 |
| Model 2 | sex x HR-neg x gamma | 0.08 |
|  | sex x HR-ASD x gamma | 0.414 |
| Model 3 | sex x HR-neg x gamma | 0.22 |
|  | sex x HR-ASD x gamma | 0.68 |

| MSEL Expressive Language T-Score vs Frontal Gamma |  |  |  |
| --- | --- | --- | --- |
|  | Slope (95% CI) | Unadjusted P Value | Adjusted P Value |
| **Model 1** |  |  |  |
| Low Risk | -13.3 (-42.1 to 15.4) | 0.359 | 0.359 |
| Males | -35.2 (-81.4 to 11.0) | 0.13 | 0.17 |
| Females | 12.4 (-19.4 to 44.2) | 0.43 | 0.43 |
| High Risk | -25.5(-43.9 to -7.03) | 0.007 | 0.014 |
| Males | -24.1(-51.5 to 3.2) | 0.08 | 0.160 |
| Females | -27.1 (-50.9 to -3.3) | 0.03 | 0.12 |
| **Model 2** |  |  |  |
| LR-Negative | -13.3 (-42.3 to 15.7) | 0.36 | 0.36 |
| Males | -34.4 (-81.0 to 12.3) | 0.15 | 0.30 |
| Females | 11.4 (-20.7 to 43.6) | 0.48 | 0.576 |
| HR-Negative | -14.3 (-37.4 to 8.7) | 0.22 | 0.33 |
| Males | -4.5 (-43.3 to 34.3) | 0.82 | 0.820 |
| Females | -23.6 (-49.9 to 2.7) | 0.08 | 0.24 |
| HR-ASD | -40.1 (-77.6 to -2.6) | 0.04 | 0.12 |
| Males | -41.2 (-87.7 to 5.2) | 0.08 | 0.24 |
| Females | -37.1 (-103.7 to 29.4) | 0.27 | 0.41 |
| **Model 3 (36 month Expressive T-Score)** |  |  |  |
| LR-Negative | 15.8 (-15.4 to 47.1) | 0.31 | 0.47 |
| Males | -4.4 (-52.3 to 43.6) | 0.86 | 0.86 |
| Females | 40.7 (6.3 to 75.0) | 0.02 | 0.12 |
| HR-Negative | 6.3 (-16.8 to 29.4) | 0.59 | 0.59 |
| Males | 6.2 (-28.9 to 41.2) | 0.73 | 0.86 |
| Females | 6.5 (-22.0 to 35.0) | 0.65 | 0.86 |
| HR-ASD | -33.1 (-70.0 to 0.82) | 0.06 | 0.18 |
| Males | -42.5 (-88.0 to 3.0) | 0.07 | 0.21 |
| Females | -17.3 (-74.4 to 39.7) | 0.54 | 0.86 |

† Results above are from linear regression models in which the outcome variable was MSEL Expressive Language T-Scores. The independent variables were frontal gamma power, sex, and risk (Model 1) or group (Models 2 and 3). Full factor interactions of independent variables were included in the models. Potential confounders - head circumference and maternal education, were included as covariates. Slopes presented are for Frontal Gamma Power and MSEL Expressive T-Score. Both unadjusted and adjusted *P* Values for multiple comparisons, using False Discovery Rate are presented.

##### Model 1

Slope comparisons of subgroups from Model 1 revealed that high-risk toddlers showed a significant negative effect of frontal gamma power on expressive language T-scores (unadjusted p=0.007; adjusted p=0.014), while low risk toddlers did not. However, the effect of frontal gamma power on MSEL Expressive T-Scores was not significantly different between risk groups. Risk groups were further subdivided by sex to evaluate whether the effect of frontal gamma on MSEL Expressive Language T-scores was similar between males and females. There was no significant difference between males and females.

##### Model 2

Slopes of MSEL Expressive T-scores versus Frontal Gamma Power from Model 2 are also shown in Table 3. However given the small number of participants in HR-ASD group, these results should be interpreted with caution. Between outcome groups, HR-ASD toddlers had the strongest negative association (unadjusted p=0.04, adjusted p=0.12). However, the effect was not significantly different from LR or HR-NoASD groups. When groups were further subdivided by sex, the strongest negative relationship between frontal gamma and expressive language were observed in HR-NoASD Females and HR-ASD Males (Table 3, See Supplemental Figure 2 for scatterplots).

#### Frontal Gamma and Future MSEL Language Scores

Finally we assessed associations between frontal gamma power at 24 months and later language ability at 36 months (Table 3). In this third model (Adjusted R^2^ = 0.19), LR and HR-ASD groups significantly differed in their associations (F_1.53_ =4.36, *p*=0.04), with the LR group having a positive association, and the HR-ASD having a negative association. When groups were subdivided by sex, a stronger positive association between frontal gamma power and later language scores was observed in LR females (slope 40.7, p=0.02, CI 6.3 to 75.0) compared to LR males (slope −4.35, p=0.86, CI −52.3 to 43.6).

## DISCUSSION

Here we report that at 24 months of age, resting frontal gamma power was significantly reduced in high-risk toddlers *without* ASD compared to low-risk controls; however, no difference was observed between high-risk toddlers *with* ASD and low-risk controls, suggesting that the single measure of resting gamma power is not a useful biomarker of ASD - at least at 24 months. Furthermore, higher gamma power in the high-risk group was only marginally associated with ASD outcome (p=0.06), and this association was not maintained when language ability was added as a covariate, emphasizing the strong linkage between ASD diagnosis and language skills.

Our lab’s previous longitudinal analysis from a smaller subset of this study population found that high-risk infants (collapsed across ASD outcome) at 6 months had lower power across all frequency bands, but by 24 months gamma power was similar between high- and low-risk infants(13). Our new finding that HR-NoASD toddlers have reduced frontal gamma power at 24 months supports a role for early neural compensatory mechanisms impacting ASD outcome. One possibility is that *maintenance* of reduced frontal gamma across the first two years may be a marker of improved developmental outcome.

In support of this hypothesis, low frontal gamma power was associated with better language ability in the high-risk toddlers. However, there was no such association in the low-risk group. Interestingly, in a similar age group, Benasich et al. have reported reduced frontal gamma in toddlers with familial risk for language impairment. However they did not evaluate the association between gamma and language function within this subset of children, rather they found gamma to be positively correlated with language *across* a larger sample which combined participants both with and without familial risk of language impairment (17,18). In our study we only observed this positive relationship in LR females when comparing frontal gamma power at 24 months to MSEL Expressive Language scores at 36 months. While Benasich et al. had similar numbers of males and females in their enrolled population, only a subset had EEG and behavioral data, and the breakdown of males versus females for each age group analyzed was not reported. Our data suggests that sex may play an important role in this relationship.

### Gamma and Language

Why would reduced gamma in a high-risk population be associated with improved language ability? Gamma oscillations are associated with a variety of higher order cognitive processes including language(16,51), attention(52,53), and working memory(54,55). However, gamma oscillations also indirectly represent the balance between excitatory and inhibitory neurons. Gamma oscillations in the cortex are generated by parvalbumin (PV) inhibitory interneurons, however disruption in PV interneurons in rodents has been shown to both increase and decrease spontaneous gamma power(56). Decreased gamma oscillations in the context of aberrant neurocircuitry may represent a variety of functions including successful compensation for processes that may increase gamma oscillations such as PV hypofunction. Alternatively, increased gamma in already abnormal neurocircuitry may lead to a ceiling effect, preventing further increase in gamma during cognitive processes. In this case, reduced gamma power would provide a more pliable system for learning. Teasing this out further is a challenging task. Longitudinal analysis of baseline gamma focused on differences between both group, and sex within group, will be useful. In addition, future studies evaluating the relationship between baseline gamma and evoked gamma within outcome groups, and how this relates to language will improve our understanding of the developmental role of gamma oscillation within high-risk populations.

### Sex Differences

Given the growing evidence of sex differences in early brain development and plasticity in ASD(57–61), in addition to differences in prevalence and phenotype between sexes, this study closely examined any possible within-group sex differences. Prospective studies of familial high-risk infants provide a unique opportunity to investigate possible compensatory mechanisms that “protect” females from ASD. Given our limited sample size, strong conclusions cannot be made with regard to sex differences. However presenting and evaluating data sub-grouped by sex is important for building hypotheses for future studies. In this study, female high-risk toddlers with ASD (n=5) had significantly lower receptive language skills than their male counterparts, and increased ADOS severity scores. Reduced IQ in females with ASD has been observed by several other groups(62,63), however others, specifically investigating high-risk infants in a larger sample size than this study, did not observe within-group sex differences in cognitive functioning or ASD symptoms severity(64). In this study, there were no significant differences between males and females across outcome groups in frontal gamma power at 24 months. However, when individual data points are examined, highrisk females make up a larger proportion of the lowest quartile of mean frontal gamma power (Figure 2). Furthermore, when the high-risk group is further separated by ASD outcome, it is the HR-NoASD females and HR-ASD males that have the strongest negative relationship between frontal gamma power and language ability. One possible explanation for this similarity is that these two subgroups have the greatest similarities in underlying neurobiology. While few studies have focused on genetic risk factors in *unaffected* high-risk females, the increased genetic burden observed in females with ASD suggests that at least a portion of unaffected high-risk females have a genetic burden similar to that seen in affected males, but do not develop ASD symptoms.

### Limitations

This study has several limitations. Given the longitudinal nature of the study, EEG acquisition changed over the course of the study. Two types of nets were utilized and EEGs were collected at two sampling rates. Given this variation we utilized batch preprocessing methods and artifact removal specific for infant EEG data to reduce any additional differences in data analysis. In addition, analyzed electrodes for each net type were carefully selected using EGI published reports(65) to ensure the same regions of interest were represented for each net type. A second limitation is that while this was a large study, enrolling over 100 HR infants, our sample size of HR-ASD toddlers with high quality EEG data at 24 months was small (n = 16), limiting our statistical power within this group. Finally, it should be noted that our participants, including those diagnosed with ASD, generally had age-appropriate language abilities. Limited variability of language skills within groups may have hindered our ability to observe statistically significant associations.

### Conclusions

We found that high-risk toddlers *without* ASD have reduced baseline frontal gamma activity, and that within this study’s high-risk population low frontal gamma power was associated with better language ability. Furthermore, this negative association between gamma power and language was largely driven by the high-risk females, emphasizing the importance of sex subgroup analysis. Together these findings suggest that gamma oscillations at this age may represent the result of ongoing compensatory mechanisms. To better understand the role of gamma oscillations in ASD, we must disentangle longitudinal compensatory changes in neural circuitry from core features of brain dysfunction. This requires both longitudinal analysis of high-risk populations, starting very early in life, as well as continued investigation into the relationship between baseline and evoked gamma oscillations throughout the course of early development.

## ABBREVIATIONS

ADHD: : Attention Deficit Hyperactivity Disorder
ADOS: : Autism Diagnostic Observation Schedule
ASD: : Autism Spectrum Disorder
BEAPP: : Batch EEG Automated Processing Platform
EEG: : Electroencephalography
HAPPE: : Harvard Automated Preprocessing Pipeline for EEG
HR-ASD: : High-risk with ASD
HR-NoASD: : High-risk without ASD
LR: : Low-risk without ASD
MARA: : Multiple Artifact Rejection Algorithm
MSEL: : Mullen Scales of Early Learning
PDDST-II: : Pervasive Developmental Disorders Screening Test-II PV: parvalbumin
SCQ: : Social Communication Questionnaire

**Author Contributions:**
CLW was involved in study conception, performed the EEG and behavioral data analysis, interpreted the data, and drafted the manuscript. ARL and LJGD contributed to EEG preparation and analysis, and critically revised the manuscript for intellectual content. HTF and CAN were responsible for the study design, overseeing data acquisition, and critically reviewed the manuscript for intellectual content.

## Acknowledgements

We thank all the families and staff who were involved in this study. We also thank Adriana Sofia Mendez Leal and Michael Mariscal for contributing code used in EEG analysis.

This research was Support for this work was provided by: The National Institutes of Health (R01-DC010290 to HTF and CAN; R21 DC 08637 to HTF; 1T32MH112510 to CLW), Autism Speaks (1323 to HTF), Simons Foundation (137186 to CAN), FRAXA Research Foundation (CLW), American Brain Foundation (ARL), Autism Science Foundation (ARL, CLW) and Nancy Lurie Marks Family Foundation (ARL). Statistical analysis was conducted with support from Harvard Catalyst | The Harvard Clinical and Translational Science Center (National Center for Advancing Translational Sciences, National Institutes of Health Award UL1 TR001102) and financial contributions from Harvard University and its affiliated academic healthcare centers. The content is solely the responsibility of the authors and does not necessarily represent the official views of Harvard Catalyst, Harvard University and its affiliated academic healthcare centers, or the National Institutes of Health. The above funding bodies did not have any role in the design, collection, analyses, or interpretation of the data or in writing this manuscript.

## Notes

**Conflict of Interest Statement:** The authors declare that they have no conflicts of interest

